# Targeting C2 reduces ischemia-reperfusion injury-induced complement activation in preclinical human models

**DOI:** 10.1101/2025.10.27.684821

**Authors:** Elisabeth de Zeeuw, Laura M. Tool, Daniëlle Krijgsman, Eline Haspeslagh, Inge Van de Walle, Jolien Delaere, C. Erik Hack, Jeanette H.W. Leusen, Kevin Budding, Linda Gijzen

## Abstract

Kidney transplantation (KTx) is a main treatment option of end stage renal disease. KTx outcome is hampered by various factors including ischemia-reperfusion (IR) injury (IRI). Animal models suggest a role for natural IgM recognizing neoepitopes exposed on ischemic cells as a main trigger for IRI-induced complement activation. However, it is unclear if experimental data from these animal models can be extrapolated to human IRI.

We used *in vitro* human models for kidney IRI to evaluate complement activation. First, we compared IgM binding and complement fixation on different endothelial cell (EC) sources in a 2D culture model, using primary kidney-derived ECs, primary lung-derived ECs and human umbilical vein ECs (HUVECs). These cells were exposed to hypoxia followed by reoxygenation in presence of complement-active human serum, or serum subjected to targeted complement inhibition. Next, we validated our findings in a 3D microfluidic organ-on-a-chip model for human kidney IRI using both HUVECs and renal proximal tubule epithelial cells (RPTECs).

In the 2D IRI model, we observed increased binding of IgM and C3 fixation on different EC sources after ischemia and subsequent reoxygenation in presence of human serum. This was not detected when cells were exposed to normoxic culture conditions. These results were confirmed in the 3D culture model, where hypoxia followed by reperfusion with complement-active human serum also led to IgM binding and C3 fixation, particularly to HUVECs. Expression of ICAM-1, a key adhesion molecule linked to renal IRI pathophysiology, was increased on RPTECs after IR-induced complement activation on HUVECs. In both models, complement inhibition at the level of C2 inhibited the abovementioned effects of IR-induced complement activation.

These results suggest classical and lectin complement pathway involvement in IR-induced damage and identify C2 as a target for therapeutic strategies.

## Introduction

Kidney transplantation (KTx) is the main treatment option of end stage renal disease, though the availability of human kidney transplants is limited^1-3^. The success of KTx outcomes is hampered by various factors, including ischemia-reperfusion (IR) injury (IRI) of the transplant^4^. The pathophysiology of IRI, which occurs when blood flow to a tissue or organ is restored after a period of ischemia, is complex^5^. In the case of kidney transplantation, IRI can present as delayed graft function (DGF) in the acute stage, which may affect long-term outcomes^6^. At present, there is no definite therapy for kidney IRI^7^.

Current interventions in kidney IR focus, amongst others, on optimal transplant management and on modulation of the IR-induced inflammatory response, including activation of the complement system^5, 8, 9^. Observations in animal models suggest a role for natural IgM, which recognizes neoepitopes exposed on ischemic cells, as a key trigger for IR-induced complement activation^10, 11^. This activation may involve both the classical (CP) and lectin (LP) pathways of complement^12-15^. However, it remains unclear to what extent these experimental data from animal models can be extrapolated to human IRI^16, 17^. Furthermore, animal models come with ethical considerations and high costs. While several simplistic *in vitro* models have been developed, these usually lack (re)perfusion flow, which is a crucial part of IR, and do not include the complement system. This indicates the need to develop suitable and predictable human preclinical models.

In this study, we optimized an *in vitro* human model for kidney IR to evaluate complement activation. First, we compared IgM binding and complement fixation on different endothelial cell (EC) sources, using primary kidney-derived ECs, primary lung-derived ECs and human umbilical vein ECs (HUVECs) exposed to hypoxia. Next, we validated our findings in a 3D microfluidic organ-on-a-chip model of human kidney IR using both HUVECs and renal proximal tubule epithelial cells (RPTECs)^18^. Here, we report that IR induces complement activation in these models, triggered by deposition of IgM on ischemic cells. This activation can be inhibited by a CP/LP-specific complement inhibitor targeting C2^19^. By employing a kidney-on-a-chip model with an integrated complement system, this study represents a significant advance toward reliable, animal-free, human-relevant preclinical modeling of ischemia– reperfusion injury.

## Methods

### Standard protocol approvals

The Medical Ethical Committee of the University Medical Center Utrecht approved the collection and usage of primary kidney-derived ECs (protocol nr: 15-018) and primary lung-derived ECs (protocol nr: 06-144). Details of all materials used for the study are listed in Supplementary Table 1.

### Cell culture (2D model)

HUVEC cells (CRL-1730™, ATCC®) were cultured in endothelial cell growth basal medium-plus (Lonza) and EBM-Plus SingleQuots additives (Lonza), supplemented with 0.1 μg Primocin (Bio-Connect) at 5% CO_2_ and 37 °C. Primary lung ECs and primary kidney ECs were collected from lung or kidney graft perfusates, respectively, and cultured as previously described using Endothelial Cell Growth Medium MV 2 (Promocell)^20^. Primary ECs and HUVECs were passaged when confluency exceeded 80%. Cells were detached using Accutase cell detachment solution (Sigma-Aldrich), pelleted (450 x *g*, 5 min), and seeded into F-bottom 96-well plates (Thermo Scientific) for use in experiments.

### Hypoxia and reoxygenation (2D model)

Cells were seeded at 50 000 cells/well in flat-bottom 96-well plates and cultured for 24 hours (h). Hypoxia was induced by culturing the cells in a hypoxia chamber (Ruskinn INVIVO2 500, Baker-Ruskinn) at 0.1% O_2_ and 5% CO_2_ at 37 °C for 24h (primary ECs) or 48h (HUVEC). Normoxic control cells were cultured simultaneously under standard conditions (21% O_2_, 5% CO_2_ at 37°C). Next, the medium was discarded and replaced by veronal-buffered saline, pH 7.4 (VB; Lonza), containing 30% (v/v) serum as complement source. This serum was either pooled human complement active serum (Innovative Research), heat-inactivated (HI) serum - incubated for 30 minutes (min) at 56 °C to inactivate complement - C2-depleted human serum (Complement Technology), or IgG/IgM-depleted serum (Pel-Freez biologicals). Where indicated, serum was pre-incubated for 15 min at room temperature (RT) with an equal volume of VB containing monoclonal anti-C2 antibody ARGX-117 ^19^, an isotypic antibody control for ARGX-117 (argenx), or 30 µg/mL purified human plasma C2 (Complement Technology), and then diluted with VB to yield a final concentration of 30%. Cells were then reoxygenated by culturing at 21% O_2_ and 5% CO_2_ for 2h at 37 °C, after which they were detached with Accutase, centrifuged (450 x *g*, 5 min), and resuspended in phosphate-buffered saline, pH 7.4 (PBS; Mediaroom Medical Microbiology, UMC Utrecht). Cells were transferred to a V-bottom plate (Greiner, BioOne) and stained with biotinylated anti-human IgM (Sigma) or anti-human C3 (LSBio), followed by incubation with streptavidin-APC (eBioscience) in FACS buffer (PBS-0.1%, w/v, bovine serum albumin (BSA; Roche), 0.01% NaN_3_) for 45 min on ice in the dark. Cells were also stained with annexin V (AnV, Biolegend) and 7-aminoactinomycin D (7AAD, BD) to distinguish between living and apoptotic/necrotic cells. Cells were analyzed using flow cytometry (FACS Canto II, BD Biosciences) and FlowJo software (TreeStar).

### The microfluidic 3D co-culture as a model for human kidney IRI

RPTECs (MTOX1030, Sigma-Aldrich) and HUVECs (C2519A, Lonza) were cultured as previously described^18^. At 90% confluency, cells were washed, detached, pelleted (200 x *g*, 5 min), and seeded into the OrganoPlate. RPTECs up to passage 3 and HUVECs up to passage 4 were used in the co-culture model. The 3D microfluidic RPTEC-HUVEC co-culture was established by plating cells in the OrganoPlate® 3-Lane 40 (MIMETAS BV), as described^18^. Briefly, cells were grown in a tubular conformation in direct contact with extracellular matrix (ECM) under flow conditions. Medium was changed on days 3 and 6 of culture. Polarized confluent tubules were established on day 6 or 7 of culture. Co-cultures used in experiments were no older than 10 days.

To induce ischemia in the co-culture model, plates were removed from the rocker platform and placed under static conditions in a low-oxygen incubator (1% O_2,_ 5% CO_2_, 37 °C) for 48h. To induce reperfusion, the medium of both channels was discarded and replaced with RPTEC tox medium (basal media with tox supplement; Sigma-Aldrich) in the RPTEC channel and Cell Biologics medium supplemented with either 30% HI or 30% fresh human pooled serum in the HUVEC channel. Plates were then incubated for 24h under normoxic conditions (21% O_2,_ 5% CO_2_, 37 °C) on the rocker platform to mimic reperfusion. To evaluate the effect of ARGX-117, medium with complement-active serum in the HUVEC channel was pre-incubated with ARGX-117 for 20 min at RT before addition to the OrganoPlate. The indicated ARGX-117 concentrations refer to the amount per mL of serum-supplemented medium.

### Analysis of function and complement fixation (3D co-culture model)

At baseline, after ischemia, and after reperfusion, phase contrast microscopy was used to evaluate the morphology of the cultures using the ImageXpress XLS Micro microscope (Molecular Devices, 4x magnification). Transepithelial/endothelial electrical resistance (TEER) of the cultures in the OrganoPlate was measured with an automated multichannel impedance spectrometer (OrganoTEER, MIMETAS BV) and OrganoPlate 3-Lane-fitting electrode boards, as previously reported, and expressed in Ohm*cm^2^ (Ω*cm^2^) using OrganoTEER software^21^.

To measure IgM and C3 fixation to the cells, cultures were washed with RPTEC tox medium in the RPTEC channel (100 μL/well) and CellBiologics complete medium in the ECM and HUVEC channels (50 μL/well) for 10 min on the interval rocker platform at 37°C and were fixed with 3.7% formaldehyde (Sigma) in HBSS (Thermo Fisher) for 15 min. The cells were then stained for IgM, C3, or ICAM-1. An overview of the antibodies used can be found in Supplementary Table 2. Briefly, cultures were permeabilized for 10 min using a Triton X-100 solution (0.3%, Sigma-Aldrich), blocked, and incubated with primary antibody for 2.5 h followed by incubation with secondary antibody and nuclei stain for 30 min. All incubation steps were performed on the rocker platform at RT. Fluorescent Z-series images were acquired using the ImageXpress® Micro Confocal High-Content Imaging System (ImageXpress XLS Confocal; Molecular Devices), and maximum intensity projections were saved to quantify IgM and C3 staining on HUVECs and ICAM-1 staining on RPTECs. In control experiments non-specific IgM and C3 deposition was observed in the area where the ECM and HUVEC tubule overlap on the 2D projection (i.e. the ECM interface). Only the ECM-opposite part of the HUVEC channel was included for quantification of IgM and C3 binding. For ICAM-1 staining, the complete RPTEC channel was used as the region of interest, as non-specific binding was minimal. An automatic thresholding method using the “default” option in ImageJ, based on the IsoData algorithm, was applied to separate the area of interest from the background, resulting in a binary mask of the image. The built-in plugin “Analyze Particles” was used to determine the percentage area of positive pixels in the image using the following settings: size ≥ 20, circularity = 0.00 – 1.00, “summarize” option selected, and edges included.

### ELISA for free C2

Samples to be tested were diluted in Tris-buffered saline (Thermo Fisher) supplemented with 2 mM CaCl_2_ (Sigma-Aldrich) and incubated in MaxiSorp microtiter plates coated with ARGX-117. C2 bound to the plates was detected with biotinylated mouse anti-human C2 antibody mAb32 (argenx), which binds to a different epitope on human C2 than ARGX-117. Plates were then developed with streptavidin-HRP and TMB (SDT reagents) and analyzed with a spectrophotometer. Recombinant human C2 (ImmunoPrecise) was used as a standard. The use of ARGX-117 as a capture antibody prevents binding of C2 already complexed with ARGX-117 in a sample. Hence, the assay measures unbound, free C2.

### Data analysis and statistics

Images were processed using ImageJ (version 1.52d)^22^. Data were analyzed with Excel (Microsoft office 365 Business, version 2308) and GraphPad Prism (GraphPad Software Inc., version 10.0.2). Error bars represent the standard deviation. Statistical analysis was performed on pooled data from three independent experiments for IgM and C3 immunostaining and from a single experiment for ICAM-1. One-way ANOVA was used, followed by Dunnett’s test for multiple comparisons. If the Brown-Forsythe test indicated unequal variances, the Kruskal-Wallis test was used instead of one-way ANOVA. For ICAM-1 immunostaining, data were normalized using a y=log(y) transformation. A *P*-value of ≤ 0.05 was considered significant.

## Results

### Hypoxia and subsequent reoxygenation induce IgM binding and complement activation on different types of endothelial cells

The effects of hypoxic conditions were evaluated on three EC types: HUVECs, kidney-derived ECs from two donors, and lung-derived ECs. Intracellular ATP levels of primary lung- and kidney-derived ECs substantially decreased after 24h of exposure to 0.1% O_2_, whereas for HUVECs this required 48h (Supplementary Figure 1). Therefore, in subsequent experiments, 24h hypoxic conditions were used for primary kidney and lung ECs, and 48h for HUVECs. Reoxygenation in the presence of 30% serum of ECs previously exposed to ischemia induced both IgM binding and C3 fixation (Figure 1A, B). C3 fixation, but not IgM binding, to HUVECs and primary ECs was abolished when HI serum was used during reoxygenation. Moreover, enhanced IgM binding and C3 fixation occurred predominantly on AnV-positive, 7AAD-negative, and double-positive cells, but not on double-negative cells, suggesting that apoptosis or membrane flip-flop was associated with IgM binding and complement activation. A representative experiment with kidney EC donor 2 is shown in Figure 1C-E. Thus, hypoxia and subsequent reoxygenation in the presence of serum had similar effects on different EC types, in that it induced binding of IgM and C3 to early- and late-apoptotic cells. This was also seen with HUVECs, which were used as the EC source for subsequent 3D co-culture experiments.

**Figure 1.**
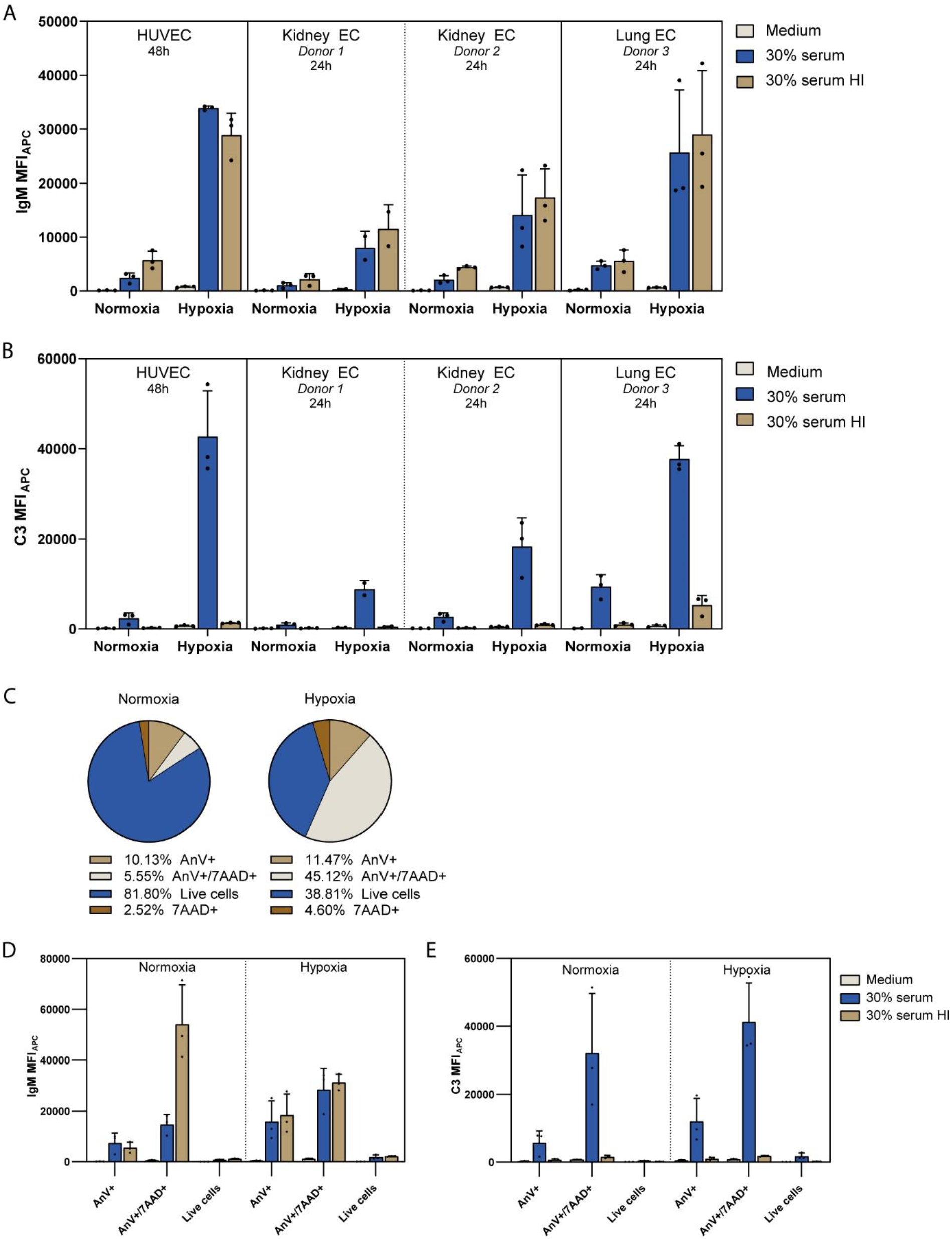
IgM binding to and complement activation on endothelial cells exposed to hypoxia and reoxygenation in the presence of fresh human serum. **A**. IgM binding to different ECs after 24/48h exposure to hypoxia/normoxia followed by 2h reoxygenation in 30% serum. **B**. C3 fixation on ECs after 24/48h culturing in hypoxic or normoxic conditions and subsequent reoxygenation with 30% serum **C**. AnV and 7AAD analysis of kidney ECs (donor 2) shows a higher percentage of AnV+/7AAD+ cells after hypoxia and reoxygenation. **D**. AnV and 7AAD profile of kidney ECs (donor 2) that bind IgM after 24h exposure to hypoxia/normoxia followed by 2h reoxygenation in 30% serum. **E**. AnV and 7AAD profile of kidney ECs that fix C3 after 24h exposure to hypoxia/normoxia followed by 2h reoxygenation in 30% serum. 7AAD: 7-Aminoactinomycin D, AnV: Annexin V, APC: allophycocyanin, h: hours, HI: heat inactivated, MFI: mean fluorescent intensity.

### 3D *in vitro* co-culture model for kidney IRI in the presence of human serum

To evaluate a potential effect of IR-induced complement activation on kidney tubule cells, a previously published 3D *in vitro* co-culture model of kidney IRI was adapted by including human serum in the HUVEC channel during the reperfusion phase^18^, as summarized in Figure 2A-C. Phase contrast images showed that addition of human serum (up to 30%) to the HUVEC channel during reperfusion under normoxic conditions (21% O_2_, perfusion flow) did not induce morphological alterations of RPTECs or HUVECs (Figure 2D). The positive control staurosporine, which induces apoptosis^23^, caused severe damage, as evidenced by rounded cells, visible holes in the tubules, and partial loss of cells in both compartments. The barrier function of the RPTECs, as assessed by measuring TEER, was slightly decreased after 24h of reperfusion with medium alone, while the barrier function of HUVECs was stable (Figure 2E), in line with the phase contrast images. Notably, TEER of both compartments was not affected by the presence of human serum during reperfusion, indicating that complement-active human serum alone did not affect barrier integrity. In contrast, exposing cultures to staurosporine resulted in TEER values around 0 Ω*cm^2^, indicating destroyed barriers. Overall, the viability of the 3D *in vitro* model under normoxic conditions was not affected by the presence of complement-active human serum in the HUVEC channel.

**Figure 2.**
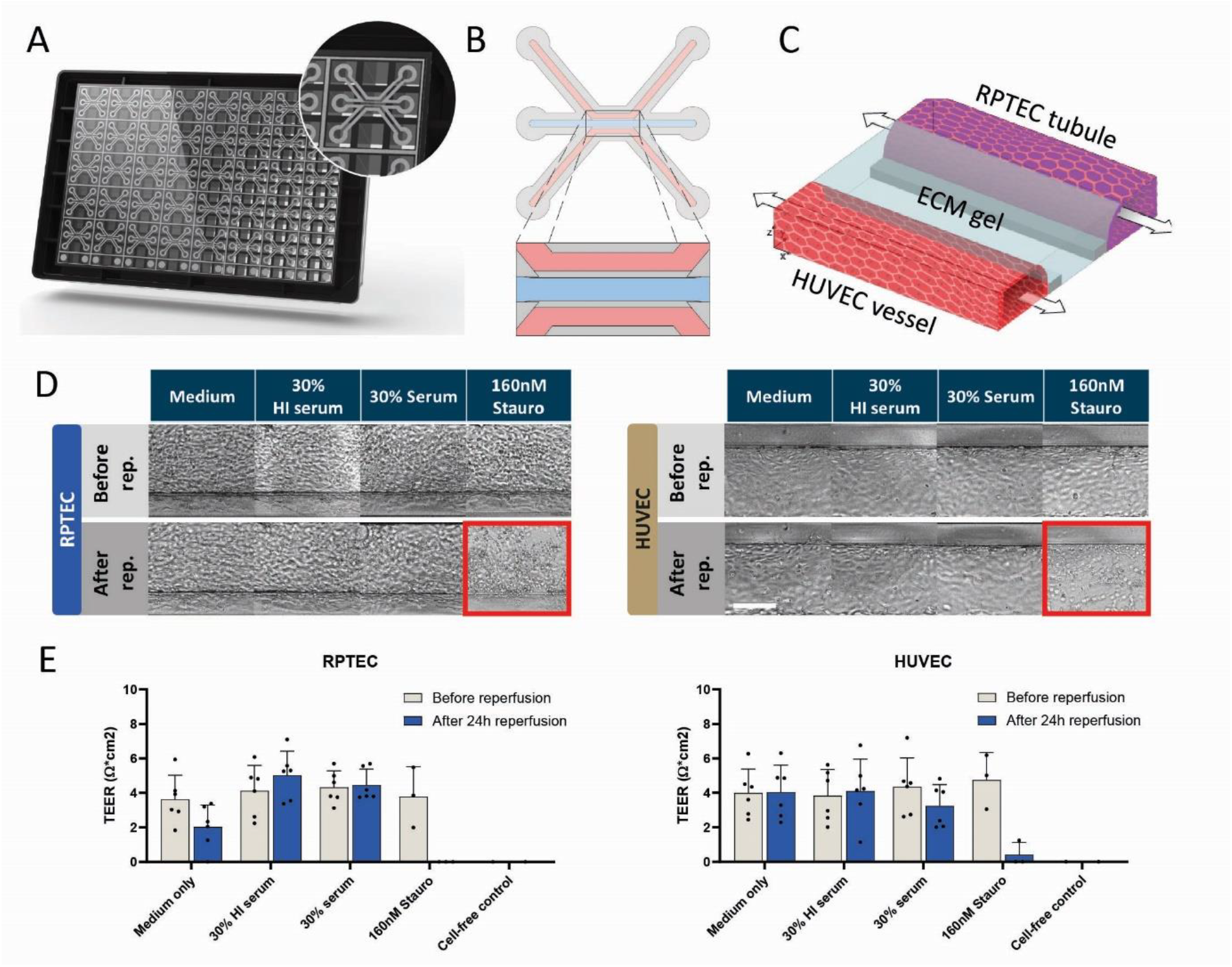
Serum compatibility and reperfusion condition in a 3D kidney co-culture model. **A**. Photograph of the microfluidic culture platform, the OrganoPlate 3-lane 40, and a zoom-in on one of the 40 chips. **B**. Schematic representation of a single chip indicating three channels that join in the center and are separated by two phase guides (grey). **C**. Three-dimensional (3D) schematic of the co-culture model comprising renal proximal tubule epithelial cells (RPTEC) in the top channel, human umbilical vein endothelial cells (HUVEC) in the bottom channel, and in the middle a collagen I extracellular matrix (ECM) gel. **D**. Phase contrast images of the co-culture model upon perfusion with complement-active human serum to the HUVEC channel for 24 h. Cultures were reperfused under normoxic conditions (21% O_2_, perfusion) for 24 h, with HUVEC culture medium supplemented with complement-active serum (= serum), heat-inactivated serum (HI serum, in which complement activation is inhibited), staurosporine (stauro), or medium-only. Staurosporine control was also added to the RPTEC channel. Phase contrast imaging revealed severe cell damage after staurosporine exposure (positive control) (red square) and no detectable alterations of cells after serum exposure. Scalebar in white = 100 µm. Rep: reperfusion. **E**. Transepithelial electrical resistance (TEER) measurements show that the presence of human serum per se did not alter the barrier integrity compared to the medium-only control. Graphs show mean ± SD. n: 6 chips for serum conditions, n: 3 for positive controls.

### IR in 3D co-culture model induces complement activation leading to RPTEC damage

To evaluate IR-induced antibody binding and complement activation, cultures were exposed to ischemia (hypoxia for 48h under static conditions) and reperfused for 24h with or without serum (Figure 3A). Ischemia increased IgM binding to HUVECs compared with normoxic conditions, both in chips supplemented with human serum or HI human serum (Figure 3B, G), whereas in the absence of serum, IgM binding was not detected, as expected. In cultures with 30% human serum as a complement source during reperfusion, IR increased C3 binding to HUVECs as compared with normoxia-reperfusion cultures. Reperfusion with HI serum did not result in increased C3 fixation. (Figure 3C, H). Under normoxic conditions, limited C3 deposition was observed, mostly at the ECM-HUVEC interface, which was excluded from quantification, as control experiments had revealed non-specific binding of C3 to the ECM matrix partially overlapping with the HUVEC channel in top-view projection images (Figure 3D-F). Both RPTECs and HUVECs showed morphological alterations after 48h of ischemia, but not after normoxia, as evidenced by rounded cells, tubule detachment, and visible holes in the tubules (Figure 3I). Reperfusion without complement-active serum further increased this damage, as visualized by an increased number and size of cell clumps, an increased number and size of holes in the tubules, and increased cell detachment. HUVECs exposed to 48h of normoxia showed similar TEER values, although with more variation, as prior to normoxia (Figure 3J). In contrast, TEER values tended to be reduced in hypoxia-exposed HUVECs but did not decline much further after reperfusion under any of the conditions. The TEER of RPTECs remained stable during 48h of normoxia and subsequent reperfusion, with a small drop in the medium-only condition. In contrast, the TEER of RPTECs exposed to ischemia was lower than before ischemia and decreased somewhat further upon reperfusion. Thus, IR induced cell damage in the RPTEC channel and to a lesser extent in the HUVEC channel. Unexpectedly, the presence of complement-active serum during reperfusion did not further influence morphology or barrier function of HUVECs or RPTECs, at least not as assessed by phase contrast imaging or TEER (Figure 3I, J).

**Figure 3.**
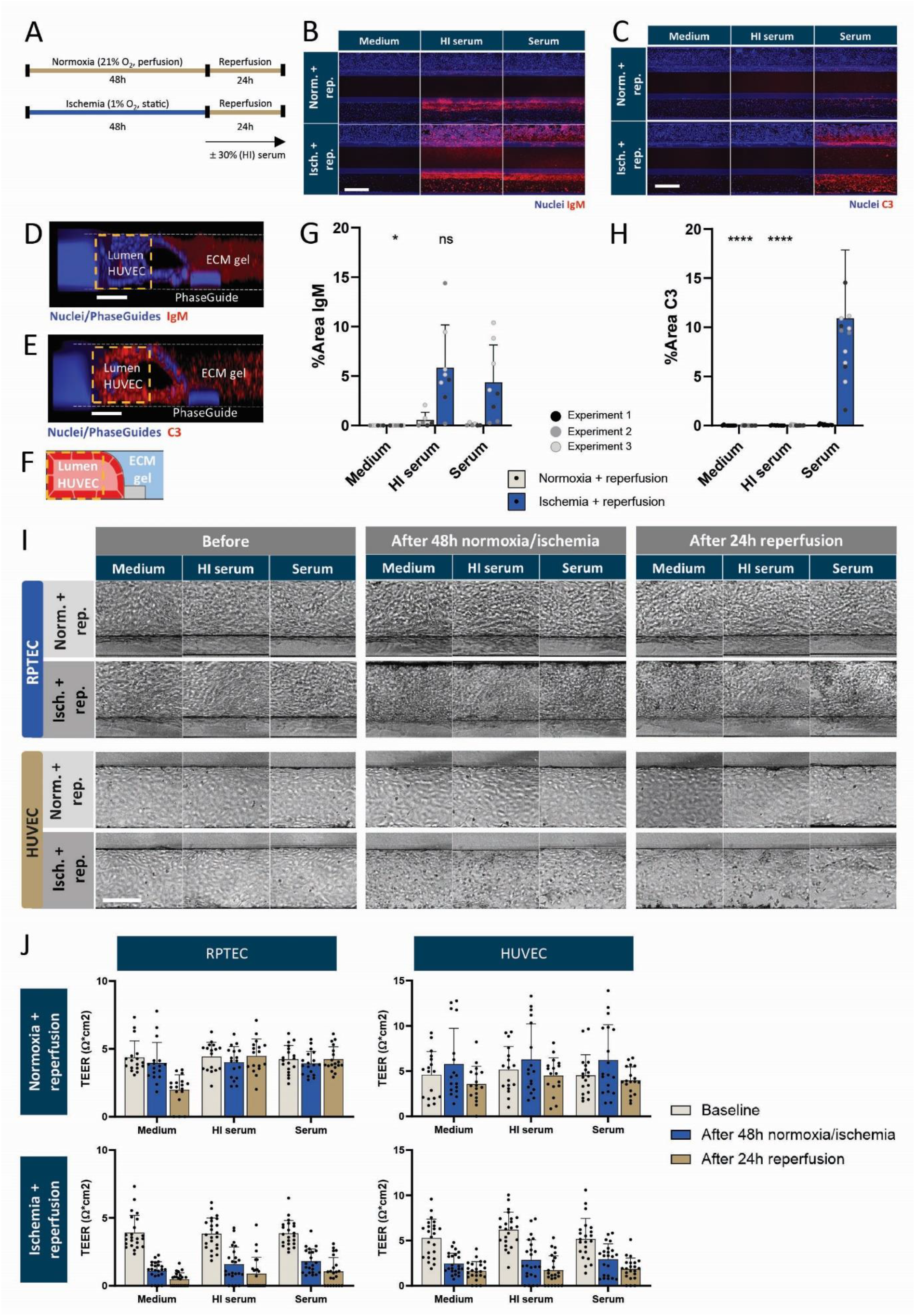
Ischemia-reperfusion induced IgM binding and complement activation in 3D co-culture model. **A**. IR was induced by 48 h exposure of the chips to ischemia (1% O_2_, static) followed by 24h reperfusion (21% O_2_, perfusion) of the HUVEC channel with medium supplemented with 30% human serum (= serum), 30% heat-inactivated serum (HI serum) or medium-only. **B**., **C**. Immunofluorescent images of IgM deposition (B, red) and C3 deposition (C, red). Note that non-specific binding of IgM in border zone of HUVEC and ECM channels in normoxia conditions. Scalebar in white = 200 µm. **D.-F**. 3D reconstruction of the HUVEC channel and ECM gel showing IgM (D, red) and C3 staining (E, red) in the ECM gel, which partially overlaps with the HUVEC channel in a top view. This overlapping area was excluded from analysis. Scalebars in white = 50 µm. **G**., **H**. Quantification of IgM (G) and C3 (H) deposition in the HUVEC channel expressed as % area of positive pixels in the image. Ns: not significant, *\* P* ≤ 0.05, *\*\*\*\* P* ≤ 0.0001, by one-way ANOVA (IgM) or Kruskal-Wallis test (C3). Note that IgM deposition is increased during IR in presence of (HI) serum as compared to normoxia, whereas C3 deposition only increased after IR in the presence of complement-active serum. **I** Representative phase contrast images show rounded cells, tubule detachment, and visible holes in the tubule after ischemia, but not after normoxia, which was exacerbated after reperfusion with medium, HI serum, or serum. Scalebar in white = 100 µm. **J**. TEER of RPTECs is decreased after ischemia (indicating disrupted barrier integrity), but not after normoxia, and remained stable after reperfusion. TEER of HUVECs only slightly decreased after ischemia as compared to normoxia and remained stable after reperfusion. Graphs show mean ± SD, and individual data points. ECM: extracellular matrix, HUVEC: human umbilical vein endothelial cells, isch: ischemia, norm: normoxia, rep: reperfusion, RPTEC = renal proximal tubule epithelial cells. n: 6-8 chips per condition within N: 3 independent experiments for IgM, n: 11-15 and N: 3 for C3, and n: 17-23 and N: 3 for phase contrast and TEER.

### Hypoxia-induced complement activation on ECs is dependent on C2

Previous studies suggest that the CP and/or LP contribute to complement activation on human cells exposed to IR^10, 24^. To elucidate the mechanism of complement activation by ischemic ECs we first used complement active IgG/IgM-depleted human serum as source of complement in the 2D culture model. When this serum was added during reoxygenation, IgM binding was abolished, as was C3 fixation (Figure 4A), indicating that complement activation in this model was IgM-induced. Next, we evaluated the effect of C2-depleted serum, with or without supplementation of physiological concentrations of C2 during reperfusion. The use of C2-depleted serum during reoxygenation did not affect IgM binding to ECs (Figure 4B). In contrast, C3 fixation was markedly reduced (Figure 4C). C3 fixation was restored by supplementing the deficient serum with purified human C2. However, C3 fixation by ischemic cells upon incubation with C2-depleted serum was not reduced to the same extent as was seen with HI serum (Figure 4C). To rule out a contribution of the alternative pathway (AP) on the observed C3 fixation, C2-depleted serum was pre-incubated with purified C2 and the anti-C2 antibody ARGX-117, which blocks C2 activation. This pretreatment of C2-depleted serum did not alter IgM binding to the ischemic ECs (Figure 4B) but reduced C3 fixation to the same extent as seen with HI serum (Figure 4C). Thus, the increased C3 fixation observed with C2-depleted serum compared to that seen with HI serum was due to the presence of trace amounts of C2 in the C2-depleted serum. Similar findings were observed with primary kidney ECs (Figure 4D, E). These results showed that complement activation by primary kidney ECs and HUVECs exposed to hypoxia and reoxygenation involved IgM and CP and/or LP, and was effectively inhibited by targeting C2.

**Figure 4.**
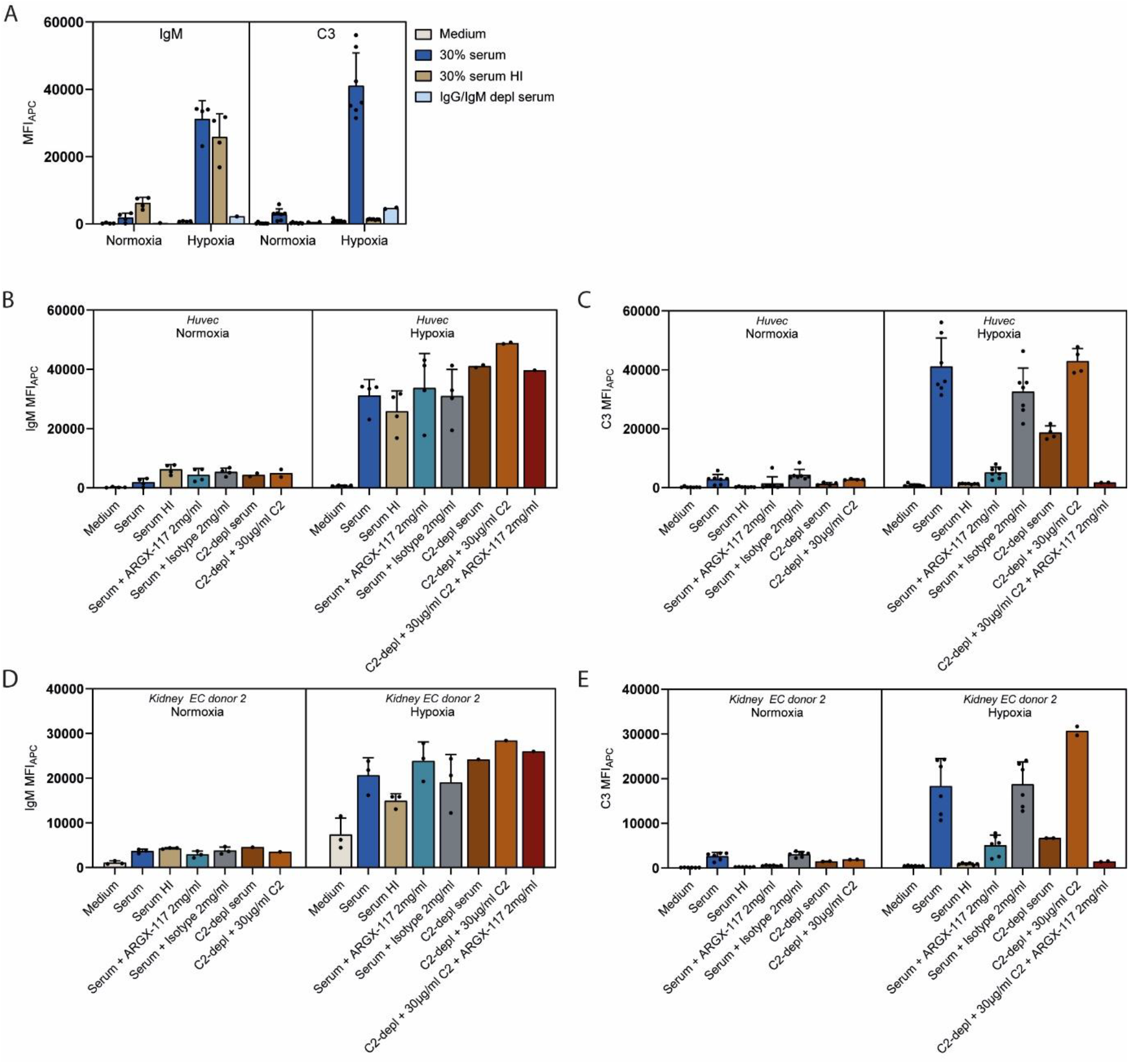
Involvement of IgM and C2 in complement activation by HUVECs and renal ECs exposed to hypoxia and reoxygenation in the presence of human serum. **A**. Reoxygenation of hypoxic HUVECs in the presence of 30% serum increased IgM binding and C3 fixation. C3 fixation was not observed after incubation with IgM/IgG depl serum. **B**. IgM binding to HUVECs after exposure to hypoxia or normoxia for 48 h followed by 2h reoxygenation in 30% serum, serum supplemented with anti-C2 antibody ARGX-117, serum depleted for C2 either or not supplemented with C2 and ARGX-117. **C**. C3 fixation of the experiment shown in B. **D**., **E**. Same experiment as in B and C, respectively, but with kidney ECs (donor 2) exposed to hypoxia/normoxia for 24 h followed by 2 h reoxygenation. APC: allophycocyanin, depl: depleted, HI: heat inactivated, h: hours, MFI: mean fluorescent intensity.

### Targeting C2 inhibits IR-induced complement activation in the 3D co-culture model

ARGX-117 was similarly tested in this co-culture model by adding different concentrations to the HUVEC channel during the reperfusion phase. In agreement with the 2D model results (Figure 4), ARGX-117 did not impact IgM deposition on IR-exposed HUVECs (Figure 5A, C), but dose-dependently decreased IR-induced C3 deposition on HUVECs (Figure 5B, D). A significant reduction in C3 deposition on HUVECs was observed after the addition of 2 mg/mL ARGX-117 (*P* = 0.016) (Figure 5D). IgM and C3 depositions were also observed on the RPTECs in some chips exposed to IR and supplemented with complement-active human serum in the HUVEC channel (Figure 5A, B). Upon addition of ARGX-117, decreased C3 deposition was observed in the RPTEC channel, most pronounced at 2 mg/mL ARGX-117 (the highest concentration) (Figure 5B). Supporting its mechanism of action, ARGX-117 dose-dependently reduced free C2 concentrations in the HUVEC channel (Figure 5E). Interestingly, free C2 was also detected in the RPTEC channel, with higher concentrations in cultures exposed to IR compared with normoxic cultures (Figure 5F). It is hypothesized that diffusion of C2 from the HUVEC channel to the RPTEC tubule is increased upon IR due to tubule damage and subsequent impaired barrier function. Analysis of the albumin levels, originating from the serum-containing HUVEC channel, in the RPTEC channel after IR supported these findings (data not shown), as did the reduced HUVEC TEER values after IR.

**Figure 5.**
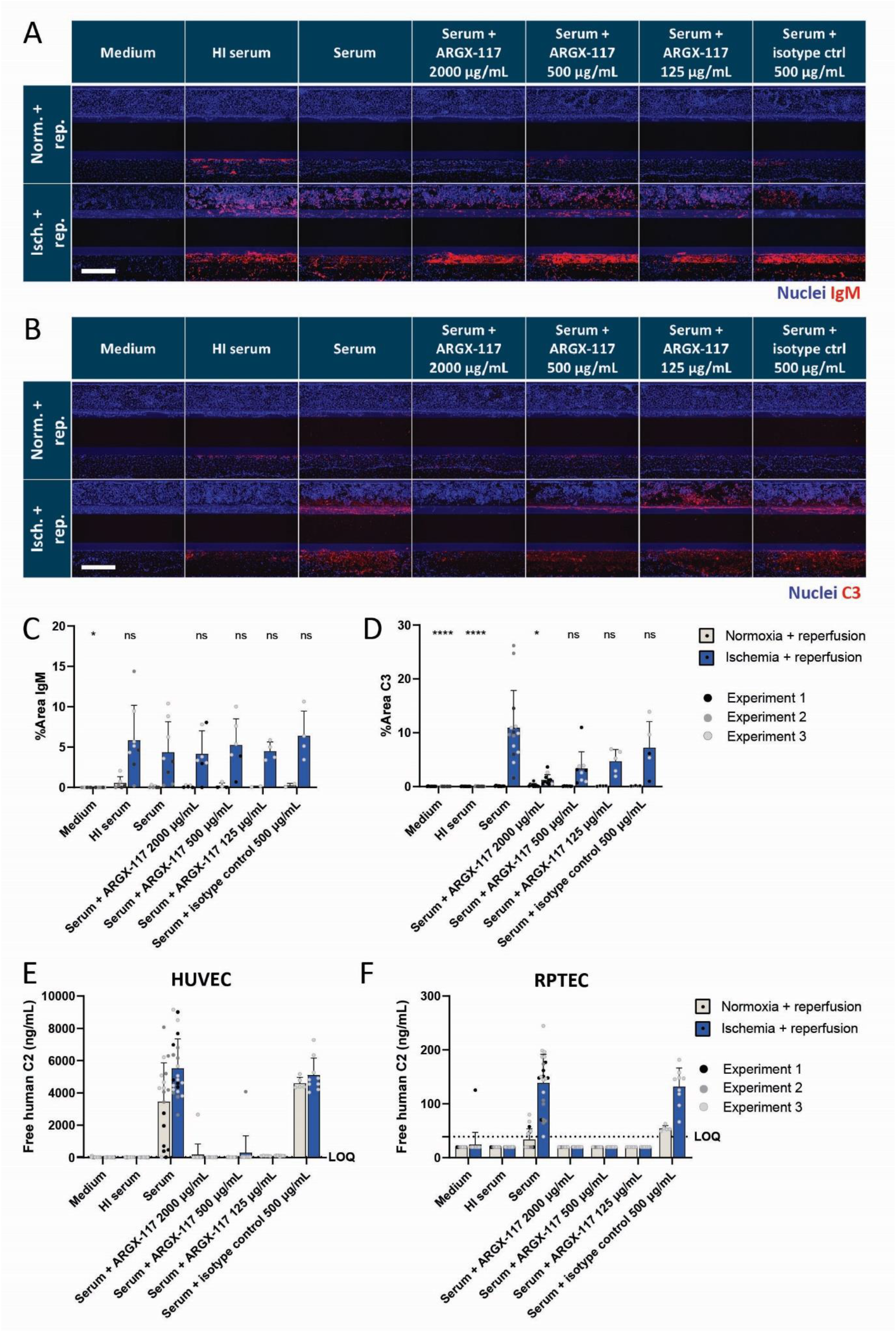
Effect of anti-C2 antibody ARGX-117 on IR-induced complement activation in the 3D kidney co-culture model. IR was induced by 48-hour exposure to ischemia (1% O2, static), or normoxia as a control, followed by 24h reperfusion (21% O2, perfusion) of the HUVEC channel with medium containing 30% human complement-active serum (= serum) with or without ARGX-117 or isotype control. **A**., **B**. Immunofluorescent images of IgM deposition (A, red) and C3 deposition (B, red). Scalebar in white = 200 µm. **C**., **D**. Quantification of IgM (C) and C3 (D) deposition in the HUVEC channel as % area of positive pixels in the image. ns = not significant, *\* P* ≤0.05, **** *P* ≤ 0.0001, one-way ANOVA (IgM) or Kruskal-Wallis test (C3). Note that IR increased IgM deposition and C3 fixation as compared to normoxia-reperfusion, and that ARGX-117 does not affect IgM binding but dose-dependently blocked IR–induced C3 deposition. **E**., **F**. Free C2 in culture supernatant was measured to determine the ability of ARGX-117 to bind C2. Supernatants of the HUVEC channel (E) and RPTEC channel (F) were tested. Free C2 was detected in culture supernatant supplemented with complement-active serum. ARGX-117 dose-dependently reduced free C2 levels. Graphs show mean ± SD of n: 2-8 chips per condition within N: 3 independent experiments for IgM, n: 3-15 and N: 3 for C3, and n: 6-22 and N: 3 for free C2. ECM: extracellular matrix, HUVEC: human umbilical vein endothelial cells, isch: ischemia, LOQ: limit of quantification, norm: normoxia, rep: reperfusion, RPTEC = renal proximal tubule epithelial cells.

### IR-induced complement activation increases ICAM-1 expression on RPTECs, which is reduced by ARGX-117

To better understand whether IR-induced complement activation could increase the inflammatory response against the kidney tubules, the expression of ICAM-1, a key adhesion molecule linked to the pathophysiology of kidney IRI^25^, was assessed on HUVEC and RPTEC tubules. ICAM-1 expression by HUVEC and RPTEC was low to absent under normoxia-reperfusion conditions but increased in the HUVEC channel after IR, independently of the presence of complement-active or HI serum (Figure 6A). In contrast, ICAM-1 upregulation by RPTECs after IR was observed only upon reperfusion with complement-active serum, but not upon reperfusion with HI serum or medium-only, indicating that this expression is stimulated by IR-induced complement activation. Specifically, ICAM-1 upregulation was observed in the RPTEC-ECM interface (Figure 6A). 3D reconstructions confirmed that the staining was specifically located on the RPTECs and not due to artificial background staining of the ECM, as seen for IgM and C3 (Figure 6C, D). Therefore, the whole RPTEC tubule was used for quantification of ICAM-1 staining. Addition of ARGX-117 during reperfusion with complement-active serum dose-dependently reduced ICAM-1 expression by RPTECs after IR exposure, indicating a potentially beneficial effect of ARGX-117 on RPTEC IRI. It should be noted that IR-induced ICAM-1 staining of the RPTEC channel was variable. However, where substantial ICAM-1 staining was observed, it was reduced by ARGX-117 (Figure 6B).

**Figure 6.**
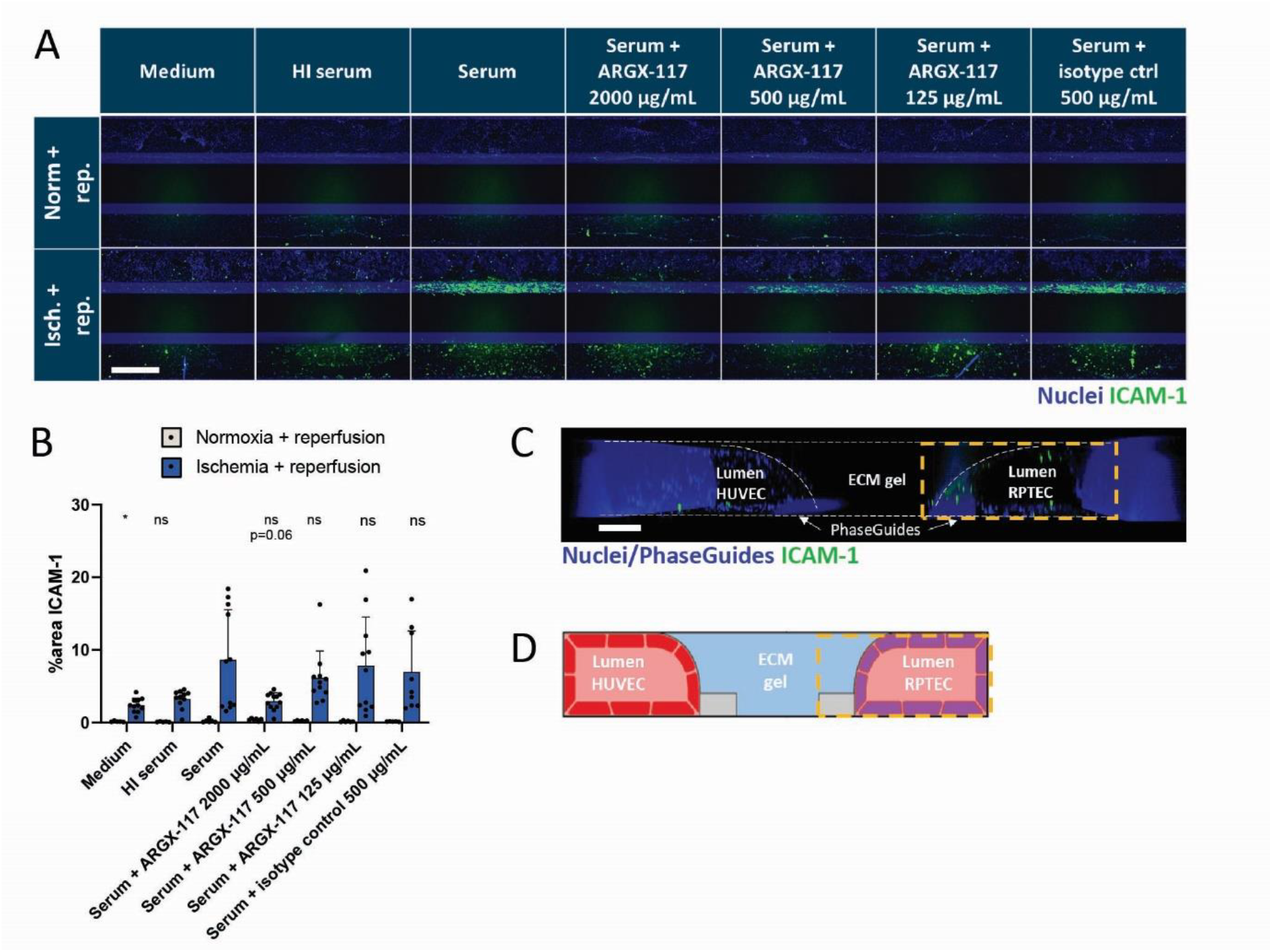
ICAM-1 expression by RPTEC is upregulated after IR in the presence of complement-active serum and inhibited by ARGX-117. Cultures were exposed to 48h ischemia (1% O_2_, static) or normoxia as control, followed by 24h reperfusion (21% O_2_) of the HUVEC channel with medium containing 30% human serum with or without ARGX-117 or isotype control, or 30% heat-inactivated (HI) serum or with medium-only. **A**. ICAM-1 is expressed (green) by ischemic RPTECs at the ECM interface upon reperfusion with medium containing complement-active serum. Addition of ARGX-117 to the complement source dose-dependently reduces IR-induced ICAM-1 expression. Scalebar in white = 200 µm. **B**. ICAM-1 expression, quantified as % area of positive pixels of the image, and expressed as mean ± SD. ns = not significant, *\* P* ≤0.05, one-way ANOVA. **C**. 3D reconstruction showed that ICAM-1 (green) was specifically stained on RPTEC cells, but not in the ECM channel, hence the complete RPTEC channel was used for quantification (orange square). Scalebar in white = 50 µm. **D**. Schematic image showing the side view of the co-culture as in the 3D reconstruction. ICAM-1 = Intercellular Adhesion Molecule 1. n: 5-12 chips per condition, N: 1.

## Discussion

Animal models have yielded a wealth of data on the role and mechanisms of complement activation in IRI^9^, but this has not yet resulted in an effective therapy for human IRI. One explanation may be that translation from animal models to the clinic is challenging and often disappointing, though studies in these models have certainly contributed to our understanding of the pathophysiologic basis of human disease^26^. One way to improve translatability is to test concepts from animal models in human *in vitro* systems, such as organs-on-chip^18^. Here, we used this approach to gain further insight into the mechanisms of complement activation in human kidney IRI. We tested various EC types in 2D cultures, including primary kidney- and lung-derived ECs as well as HUVECs. Exposure to hypoxic conditions followed by reoxygenation in presence of complement-active human serum induced binding of IgM and fixation of C3 to all three types of ECs, an effect not observed under normoxic conditions. Next, HUVECs were used that are widely available, in a 3D microfluidic IR co-culture model for human kidney IR, together with RPTECs. In this model, exposure to ischemia followed by reperfusion of the HUVEC channel with complement-active human serum also led to IgM binding and C3 fixation, particularly on HUVECs. Finally, anti-human C2 antibody ARGX-117 effectively prevented IR-induced complement activation during reoxygenation - both in primary kidney ECs in the 2D model and in HUVECs (2D and 3D).

In line with other reports, all three EC sources cultured in 2D under hypoxia showed reduced intracellular ATP levels^27^. However, primary kidney- and lung-derived ECs were more sensitive to hypoxia, showing significantly reduced intracellular ATP levels after 24h, whereas this took 48h in HUVECs. This difference may be due to adaptive mechanisms such as induction of hypoxia-associated proteins^28^ or differential gene regulation^29, 30^. Stempien-Otero *et al*. also reported minimal loss of viability of HUVECs after 24h of hypoxia, with a more substantial loss of 45% after 48h^31^. Aside from requiring longer exposure to hypoxia, HUVECs showed a similar response to reoxygenation following hypoxia as kidney or lung ECs.

Natural IgM targeting neoepitopes exposed by ischemic tissues is a key driver of complement activation during IRI in murine models^24, 32^. Use of IgG/IgM-depleted serum as a complement source during reoxygenation eliminated IgM binding and prevented C3 fixation on ischemic human ECs in the models described here, supporting a comparable mechanism of complement activation in human IRI^33^. We performed the reoxygenation experiments in presence of 30% human serum, which allows for AP activation. Our results indicate that in presence of C2 depleted serum or pooled complement-active serum pre-treated with anti-C2 antibody (inhibiting both the CP and LP but leaving the AP intact^19^), no residual complement activation was detected. This suggests that the AP could be important for amplification, but the initial trigger for complement activation is CP/LP-dependent.

With complement activation after IR as a primary objective, assessment of subsequent complement-mediated tissue damage and prevention thereof was limited in this study, although this is widely documented in the literature *in vivo*^9, 34^. For example, in the 3D co-culture model, damage to the endothelial barrier - as assessed by phase contrast imaging or TEER measurements - was seen after IR but was not increased by the addition of complement. We hypothesize that the IR exposure without complement already induced such extensive damage, that further complement-mediated damage could not be detected. In a follow up study, focusing on complement-mediated IR tissue injury rather than complement activation itself, the potential of C2 as a therapeutic target for IRI could be confirmed on a functional level. Here, it would be especially relevant to include immune cells^35^, which are known to promote inflammation and exacerbate tissue damage and transplant dysfunction^36^.

As an intermediate marker of immune-mediated IR injury, ICAM-1 expression was assessed. ICAM-1 expression by RPTECs *in vivo* is an indicator of renal damage^37^. This adhesion molecule may contribute to injury by recruiting and activating immune cells such as neutrophils and monocytes. Indeed, treatment of animals undergoing renal IR with anti-ICAM-1 antibodies or anti-sense oligonucleotides reduces kidney IRI and improves kidney function^25, 38, 39^. Interestingly, the RPTECs expressed ICAM-1 following reoxygenation in the presence of complement-active serum in the endothelial channel, but not in the presence of HI serum, which lacks a functional complement system. These results suggest that RPTECs experienced complement-dependent stress in the 3D co-culture model following reperfusion of the endothelial channel with complement-active serum. ICAM-1 expression following IR exhibited some variability but was consistently reduced by C2 targeting across all replicates. This variability may in part reflect the transient nature of ICAM-1 expression and could potentially be minimized by further optimization of the IR exposure period. As the primary objective of this study was to evaluate complement activation as an early indicator of IR injury, we selected a 24-hour reperfusion interval. It is possible, however, that downstream processes such as ICAM-1 protein upregulation require longer reperfusion times to be reliably detected.

Complement inhibitors are emerging as targeted therapies for reducing ischemic damage and subsequent DGF or rejection^40^. Eculizumab, an anti-C5 monoclonal antibody, has shown hematological and kidney improvements in patients with atypical hemolytic uremic syndrome, but has yielded mixed results in clinical trials for preventing DGF after KTx^41, 42^. This could be explained by the fact that upstream complement-activation products (e.g. the formation of the anaphylatoxin C3a) that are reported to be important drivers of IRI pathogenesis^43^ are not inhibited at the level of C5. ARGX-117 is a novel therapeutic antibody that inhibits C2^19^, thereby blocking pathologic activation of the CP and LP, as may occur in IRI^12-15^. ARGX-117 reduced complement activation in other human disease models in which complement activation is driven by IgM^19, 44^. In the 3D model described here, ARGX-117 dose-dependently decreased C3 deposition on both the endothelium and kidney epithelium when added to the endothelial channel during reperfusion. Interestingly, ARGX-117 also reduced ICAM-1 expression by HUVECs and RPTECs following ischemia and reperfusion with complement-active serum, indicating its potential to reduce complement-dependent stress on tubular epithelial cells in kidney IRI. This 3D IR model is the first step in development of an in vitro model for the study of complement-dependent ischemia reperfusion injury, with the goal to eliminate the ethical concerns, high costs and limited translatability that accompany currently used animal models. A current phase 2 study evaluating the safety, efficacy, and tolerability of ARGX-117 in deceased kidney donor transplant recipients at risk for DGF (NCT05907096) will further assess the translatability of the results described here to humans with kidney IRI.

## Data availability statement

Data from this study will not be deposited in a public repository, as no large datasets are presented. All data generated or analyzed in this study and the methods were included in the main text and the supplementary material for this article. Other source data, anonymized data, and documentation of this study will be shared upon reasonable request from any qualified researcher. Standard data sharing agreements apply.

## Declaration of generative AI and AI-assisted technologies in the writing process

The authors did not make use of any Generative AI or AI-assisted technologies during the preparation of this manuscript.

