## Supplementary document for "Targeting C2 reduces ischemia-reperfusion injury-induced complement activation in preclinical human models"

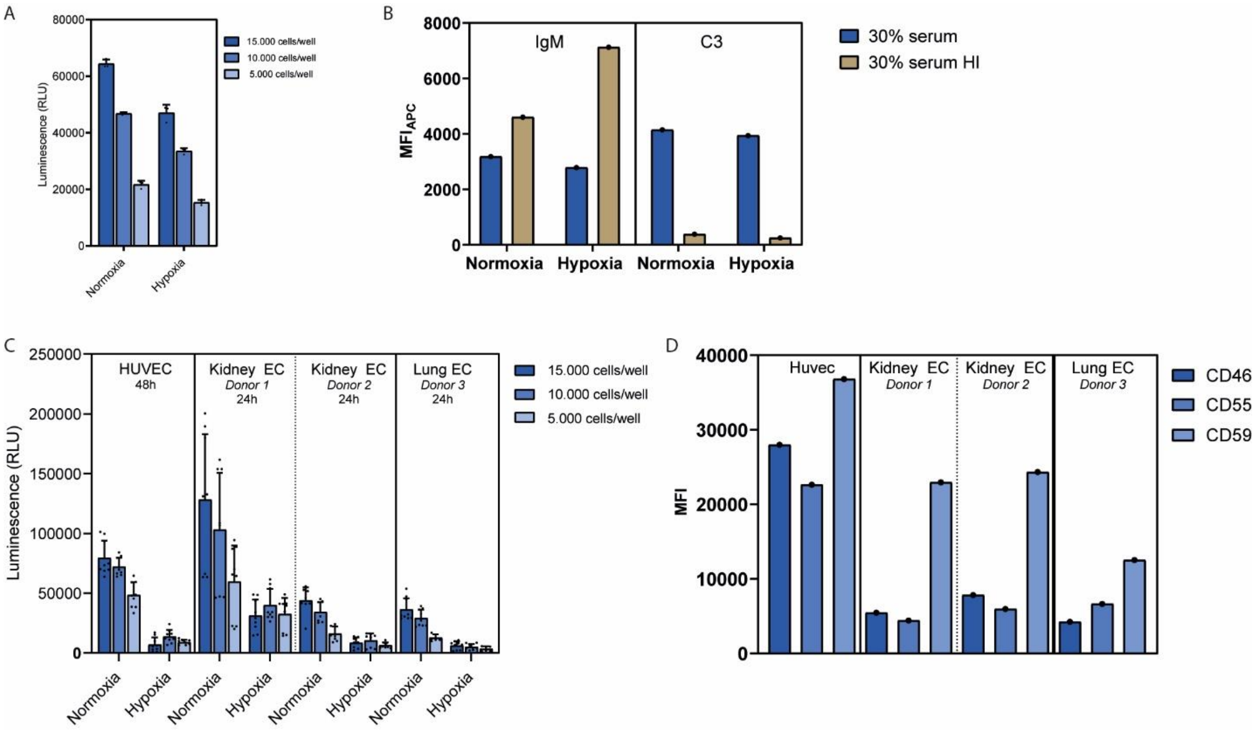

**Supplementary Figure 1. Effect of hypoxia on ATP levels in endothelial cells. A.** Exposure to 24h of hypoxia (0.1% oxygen) did not result in a large decrease in ATP levels for HUVECs. **B.** Exposure to 24h hypoxia did not result in an increase in IgM binding or C3 fixation on HUVECs. **C.** Exposure of HUVECs (48h) and primary endothelial ECS (24h) to hypoxia resulted in a decrease in cellular ATP levels compared to normoxic conditions. **D**. Expression levels of complement regulatory proteins on the surface of endothelial cells. APC: allophycocyanin, ATP: adenosine triphosphate, HI: heat inactivated, h: hours, MFI: mean fluorescent intensity, RLU: relative light units

***Supplementary Table 1** Materials used for the study*

| Material | Catalog number | Vendor |
|---|---|---|
| HUV-EC-C cells | CRL-1730 | ATCC |
| Endothelial cell growth basal medium-plus | CC-5036 | Lonza |
| EBM-Plus SingleQuots additives | CC-4542 | Lonza |
| Primocin | ant-pm-2 | Bio-Connect |
| Endothelial Cell Growth Medium MV 2 | C-22022 | Promocell |
| Accutase cell detachment solution | A6964 | Sigma-Aldrich |
| F-bottom 96-well plates | 167008 | Thermo Scientific |
| Hypoxy working chamber Ruskinn INVIVO2 500 | LP0406 | Baker-Ruskinn |
| Veronal buffer | LO 12-624E | Lonza |
| Pooled human complement active serum | ICSER1ML | Innovative Research |
| C2-depleted serum | A312 | Complement Technology |
| IgG/IgM depleted serum | 34010 | Pel-Freez biologicals |
| purified human plasma C2 | A112 | Complement Technology |
| V-bottom plate | 651101 | Greiner BioOne |
| BSA | 10735094001 | Roche |
| Annexin V | 640908 | Biolegend |
| 7-aminoactinomycin D | 559925 | BD |
| Hanks' Balanced Salt Solution | 15266355 | Gibco |
| Luminescent ATP detection kit | ab113849 | Abcam |
| Lumitrac plate | 655075 | Greiner BioOne |
| SpectraMax M3 | | Molecular devices |
| PBS | | Mediaroom Medical Microbiology UMC Utrecht |
| T75 flask – RPTEC preculture | 431464U | Corning |
| MEME alpha Modification | M4526 | Sigma-Aldrich |
| RPTEC Complete Supplement | MTOXRCSUP | Sigma-Aldrich |
| L-glutamine | G7513 | Sigma-Aldrich |
| Gentamicin | G1397 | Sigma-Aldrich |
| Amphotericin B | A2942 | Sigma-Aldrich |
| PureCol | 5005 | Advanced BioMetrix |
| Hanks' Balanced Salt Solution | 14175095 | Thermo Fisher |
| RPTECs | MTOX1030 | Sigma-Aldrich |
| HUVECs | C2519A | Lonza |
| T75 flasks – HUVEC preculture | 156499 | Thermo Fisher |
| MV2 medium supplemented with Supplement Mix Endothelial Cell Growth Medium MV2 | C-22022 | PromoCell |
| Penicillin-Streptomycin | P4333 | Sigma-Aldrich |
| Trypsin-EDTA | CC-5012 | Lonza |
| OrganoPlate® 3-Lane 40 | 4004-400B | MIMETAS BV |
| Collagen I | 3447-020-01 | AMSbio |
| HEPES | 15630-122 | Thermo Fisher |

| | | |
|---|---|---|
| $NaHCO_3$ | S5761 | Sigma-Aldrich |
| Interval rocker platform | OrganoFlow, I-OFPR-L | MIMETAS BV |
| CellBiologics complete human endothelial cell medium | H1168 | Cell Biologics |
| RPTEC Tox supplement | MTOXRTSUP | Sigma-Aldrich |
| ImageXpress XLS Micro microscope | ImageXpress XLS Micro | Molecular Devices |
| PBS | 20012068 | Thermo Fisher |
| BSA | A2153 | Sigma-Aldrich |
| Fetal bovine serum | 16140-071 | Thermo Fisher |
| Tween 20 | P9416 | Sigma-Aldrich |
| Triton X-100 | T8787 | Sigma-Aldrich |
| Tris Buffered Saline | BP24711 | Thermo Fisher |
| CaCl2 | 21115 | Sigma-Aldrich |
| s(HS)TMB | sTMB | SDT reagents |

***Supplementary Table 2*** *Antibodies used for immunofluorescent staining*

| **Antibody** | **Catalog number** | **Vendor** | **Clone** | **Conjugate** | **Dilution** |
|---|---|---|---|---|---|
| Mouse anti-human C3 | LS-C62849 | LS Bio | Monoclonal, 6C9 | Biotin | 1:100 |
| Goat anti-human IgM | B1265 | Sigma | Polyclonal | Biotin | 1:300 |
| Streptavidin APC | 17-4317-82 | Thermo Fisher | NA | APC | 1:100 |
| Mouse anti-human ICAM-1 | Ab171123 | Abcam | Monoclonal, 1A29 | NA | 1:100 |
| Goat anti-mouse IgG | A-21037 | Thermo Fisher | Polyclonal | Alexa Fluor 750 | 1:250 |
| Hoechst 33342 | H3570 | Thermo Fisher | NA | NA | 1:2000 |
